# Robust characterization of two distinct glutarate sensing transcription factors of *Pseudomonas putida* L-lysine metabolism

**DOI:** 10.1101/557751

**Authors:** Mitchell G. Thompson, Zak Costello, Niklas F. C. Hummel, Pablo Cruz-Morales, Jacquelyn M. Blake-Hedges, Rohith N. Krishna, Will Skyrud, Allison N. Pearson, Matthew R. Incha, Patrick M. Shih, Hector Garcia-Martin, Jay D. Keasling

## Abstract

A significant bottleneck in synthetic biology involves screening large genetically encoded libraries for desirable phenotypes such as chemical production. However, transcription factor-based biosensors can be leveraged to screen thousands of genetic designs for optimal chemical production in engineered microbes. In this study we characterize two glutarate sensing transcription factors (CsiR and GcdR) from *Pseudomonas putida*. The genomic contexts of *csiR* homologs were analyzed and their DNA binding sites were bioinformatically predicted. Both CsiR and GcdR were purified and shown to bind upstream of their coding sequencing *in vitro*. CsiR was shown to dissociate from DNA *in vitro* when exogenous glutarate was added, confirming that it acts as a genetic repressor. Both transcription factors and cognate promoters were then cloned into broad host range vectors to create two glutarate biosensors. Their respective sensing performance features were characterized, and more sensitive derivatives of the GcdR biosensor were created by manipulating the expression of the transcription factor. Sensor vectors were then reintroduced into *P. putida* and evaluated for their ability to respond to glutarate and various lysine metabolites. Additionally, we developed a novel mathematical approach to describe the usable range of detection for genetically encoded biosensors, which may be broadly useful in future efforts to better characterize biosensor performance.

## INTRODUCTION

A rate limiting step in the design-build-test-learn cycle is often the test phase, wherein hundreds or thousands of genetic designs need to be evaluated for their productivity ^1, 2^. Though recent advances in analytical chemistry have dramatically increased sample throughput ^3^, transcription factor-based biosensors still offer multiple advantages over traditional chromatographic and mass-spectrometry based detection methods ^4, 5^. One of the most attractive benefits is the ability to rapidly screen constructs for the production of the target compounds via either plate-based or flow-cytometry-based assays ^1–3^, which increases throughput by orders of magnitude compared to mass-spectrometry based methods. Additionally, biosensors may offer unmatched sensitivity towards specific ligands, with some sensors having picomolar affinity ^3^. The evolution of diverse microbial metabolism has provided researchers with the ability to sense a wide array of ligands, ranging from complex natural products ^6, 7^ to small central metabolites ^1, 8^.

Diacids, polyamines, and lactams are petrochemical derivatives used to produce various polyester and nylon fibers ^9, 10^. In an effort to make production of these chemicals sustainable, many groups have developed engineered microbes to synthesize these precursors ^11–14^. The L-lysine metabolism of *Pseudomonas putida* has been leveraged both in the native host and heterologously to produce valerolactam ^15^ the diacid glutarate ^16, 17^. Recently, this utility has inspired much work to uncover missing steps in the lysine catabolism of *P. putida*. These missing steps included the discovery that glutarate is not only catabolized through the previously known coA-dependent route to acetyl-coA, but is also catabolized through a coA-independent route to succinate ^16, 18^ (Figure 1). Recent work has also demonstrated that both glutarate catabolic pathways are highly upregulated in the presence of glutarate ^18^. The *Pseudomonas aeruginosa* homolog of the ketogenic pathway regulator (GcdR) ^19^and the *Escherichia coli homolog of the* glucogenic pathway regulator (CsiR) have both been characterized. Furthermore a rigorous investigation of the P. putida homologs has now been reported ^20^..

**Figure 1:**
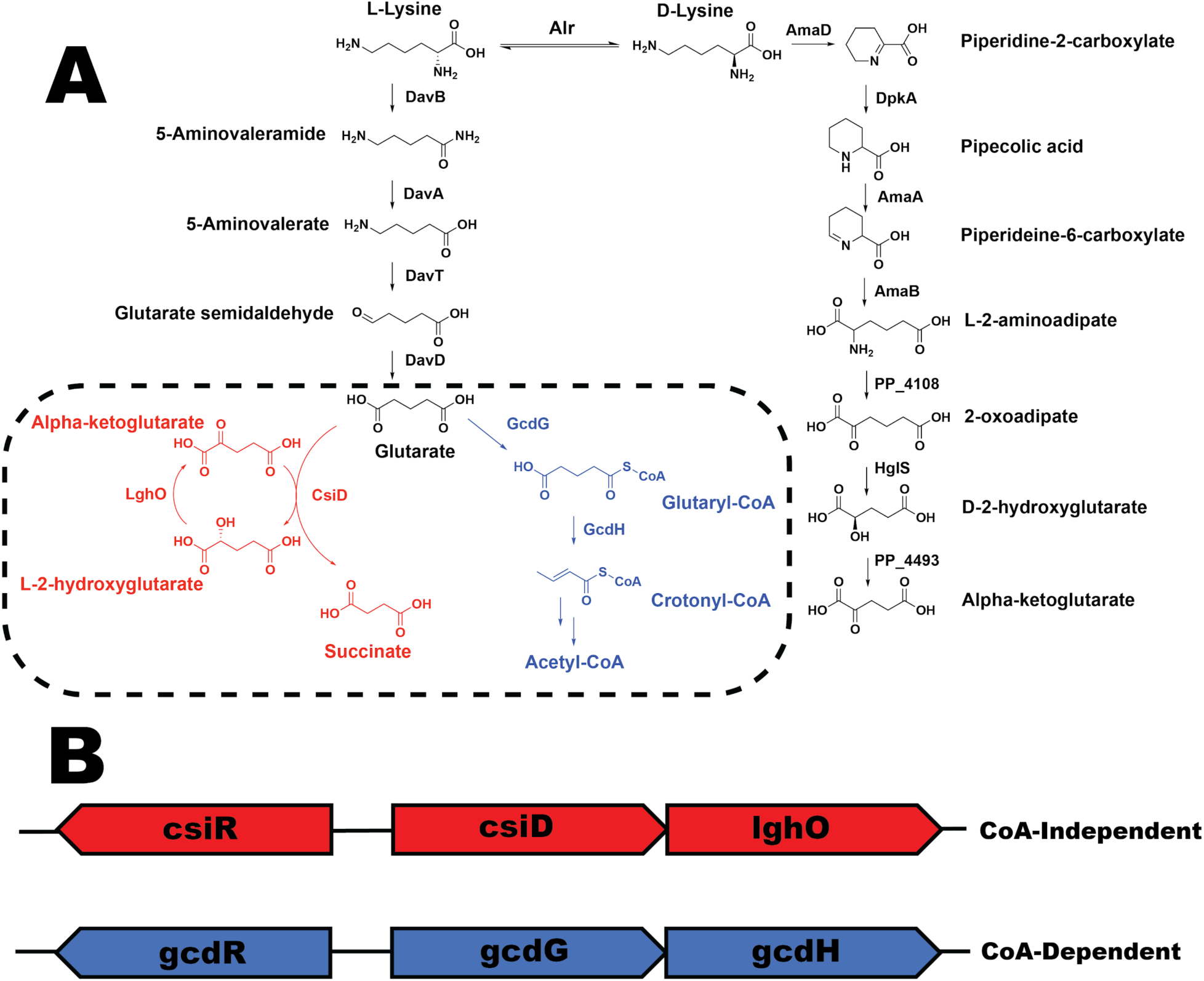
A) The known lysine catabolism of *P. putida*. Dashed box shows the two known pathways of glutarate catabolism in *P. putida*. Highlighted in red is the CoA-independent route of glutarate catabolism, in blue the CoA-dependent route. Enzymes in the metabolism are DavB - L-lysine monooxygenase, DavA - 5-aminopentanamidase, DavT - 5-aminovalerate aminotransferase, DavD - glutaric semialdehyde dehydrogenase, CsiD - glutarate hydroxylase, LghO - L-2-hydroxyglutarate oxidase, GcdG glutaryl-coA transferase, GcdH - glutaryl-CoA dehydrogenase, Alr - alanine racemase, AmaD - D-lysine oxidase, DpkA - Δ1-piperideine-2-carboxylate reductase, AmaB - pipecolate oxidase, AmaA - L-aminoadipate-semialdehyde dehydrogenase, PP_4108 - L-2-aminoadipate aminotransferase, HglS - D-2-hydroxyglutarate synthase, PP_4493 - D-2-hydroxyglutarate dehydrogenase. B) Operonic structure of the two routes of glutarate metabolism in *P. putida*.

In this work we sought to also characterize the two putative local regulators of glutarate catabolism in *P. putida*, *csiR* and *gcdR*. First, we compared the genomic context of *csiR* homologs across bacteria to bioinformatically predict a conserved DNA binding site. We then biochemically and genetically characterized both regulators. Secondly, we developed a novel mathematical approach to rigorously determine the detection ranges for genetically encoded biosensors that can be used to systematically compare biosensors. Finally, we introduced RFP transcriptional-fusions of the promoter for both catabolic pathways into *P. putida* and evaluated their induction upon the addition of various lysine metabolites.

## RESULTS

### Genomic contexts of *csiR* and *gcdR* homologs and prediction of *P. putida* binding sites

Work in *Pseudomonas aeruginosa* has characterized the GcdR regulation of ketogenic glutarate metabolism, and shown that the binding site is conserved across multiple bacterial species ^19^. While binding sites for CsiR in *E. coli* have been identified, it was yet to be investigated whether there is a conserved binding site for homologs across bacterial species ^21^. In order to identify conserved binding sites of CsiR homologs, we compared the syntenic genomic contexts of 12 selected genomes that contained neighboring *csiD* and *csiR* homologs. Genes encoding *csiR* were found in two distinct genomic contexts, either transcribed divergently from *csiD* as found in *P. putida*, or transcribed as the last gene in the *csiD* operon as in *E. coli* (Figure 2). The genomic regions upstream of the *csiD* homolog were extracted and Multiple EM for Motif Elicitation (MEME) was used to identify a conserved CsiR binding motif ^22^. A consensus A(A/G)AAATCTAGA(C/T)ATTTT motif was identified upstream of each *csiD* homolog. Previously, footprinting assays in *E. coli* BW25113 revealed two primary and two secondary bindings sites with the sequences TTGTN_5_TTTT and ATGTN_5_TTTT respectively ^21^. While this manuscript was under review Zhang et al. demonstrated via DNase I footprinting that CsiR does indeed binding at two locations upstream of *csiD*, including the conserved A(A/G)AAATCTAGA(C/T)ATTTT motif ^20^. Our consensus motif agrees closely with the secondary binding site identified, and highly suggests that the binding site of CsiR is conserved across the bacteria where homologs are present.

**Figure 2:**
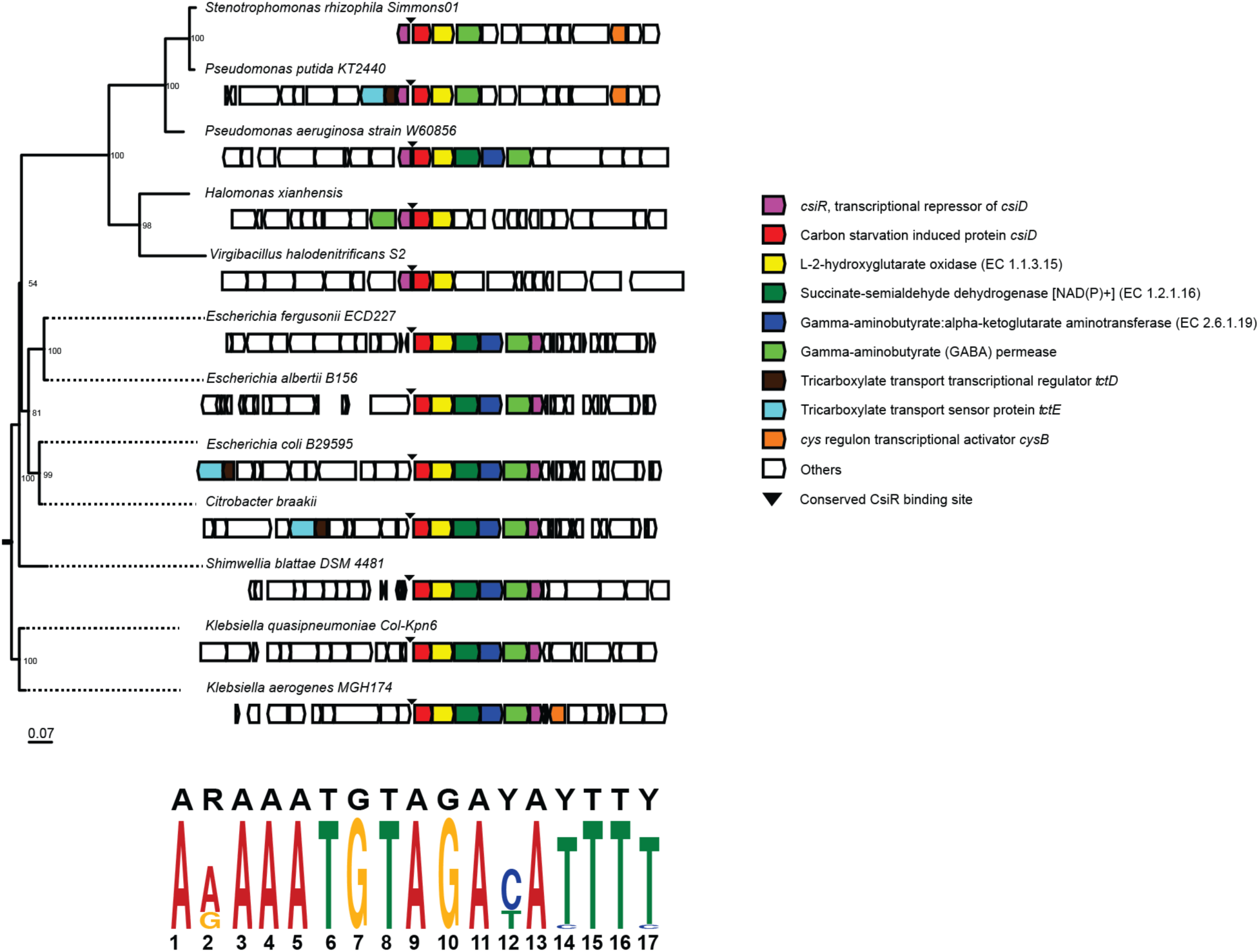
Genomic contexts of *csiR* homologs and predicted binding regions: Phylogeny of assorted gammaproteobacteria and the location of their *csiD* operons. MEME analysis of the intergenic regions upstream of the *csiD* operon resulted in the sequence motifs depicted below the tree.

### Biochemical characterization of CsiR and GcdR

To determine if the *P. putida* CsiR also acts as a regulator and to identify its putative binding sites, we biochemically characterized this protein using electrophoretic mobility shift assays (EMSAs). The CsiR protein was purified (Figure S1) and incubated with DNA probes consisting of the intergenic region between *csiR* and *csiD*. The assay showed multiple binding sites in this intergenic region, as four distinct bands appeared. These results appear to confirm previous work in *E. coli in which* four binding sites of the *E. coli* CsiR homolog were observed^21^. CsiR had a high affinity for the DNA probe, with a calculated K_d_ of approximately 30 nM (Figure 3Aa), which is similar to the 10 nM CsiR/DNA K_d_ of the *E. coli* CsiR homolog, and nearly identical to the results of Zhang et al. which showed complete probe shift at 6-fold molar excess CsiR ^20, 21^. Purified GcdR (Figure S1) also bound to its cognate probe with a single distinct shift and had an an estimated K_d_ of approximately 62.5 nM (Figure 3A) ^19^. Again, these results were consistent with Zhang et al., who showed near complete probe shifts at 8-fold molar excess GcdR ^20^. As CsiR is a GntR family transcriptional regulator, many of which act as repressors ^23^, we evaluated whether glutarate would decrease the DNA binding affinity of CsiR. EMSA assays were repeated in the presence of increasing glutarate concentrations. Analysis by gel electrophoresis revealed increasing quantities of free probe as glutarate concentrations increased (Figure 3B). These results suggest that the *P. putida* CsiR is a glutarate-responsive repressor of the *csiDlhgO* operon^21^. Zhang et al. also showed that in addition to dissociating from the DNA probe at high concentrations of glutarate, CsiR was also responsive to 2-hydroxyglutarate (2HG) ^20^.

**Figure 3:**
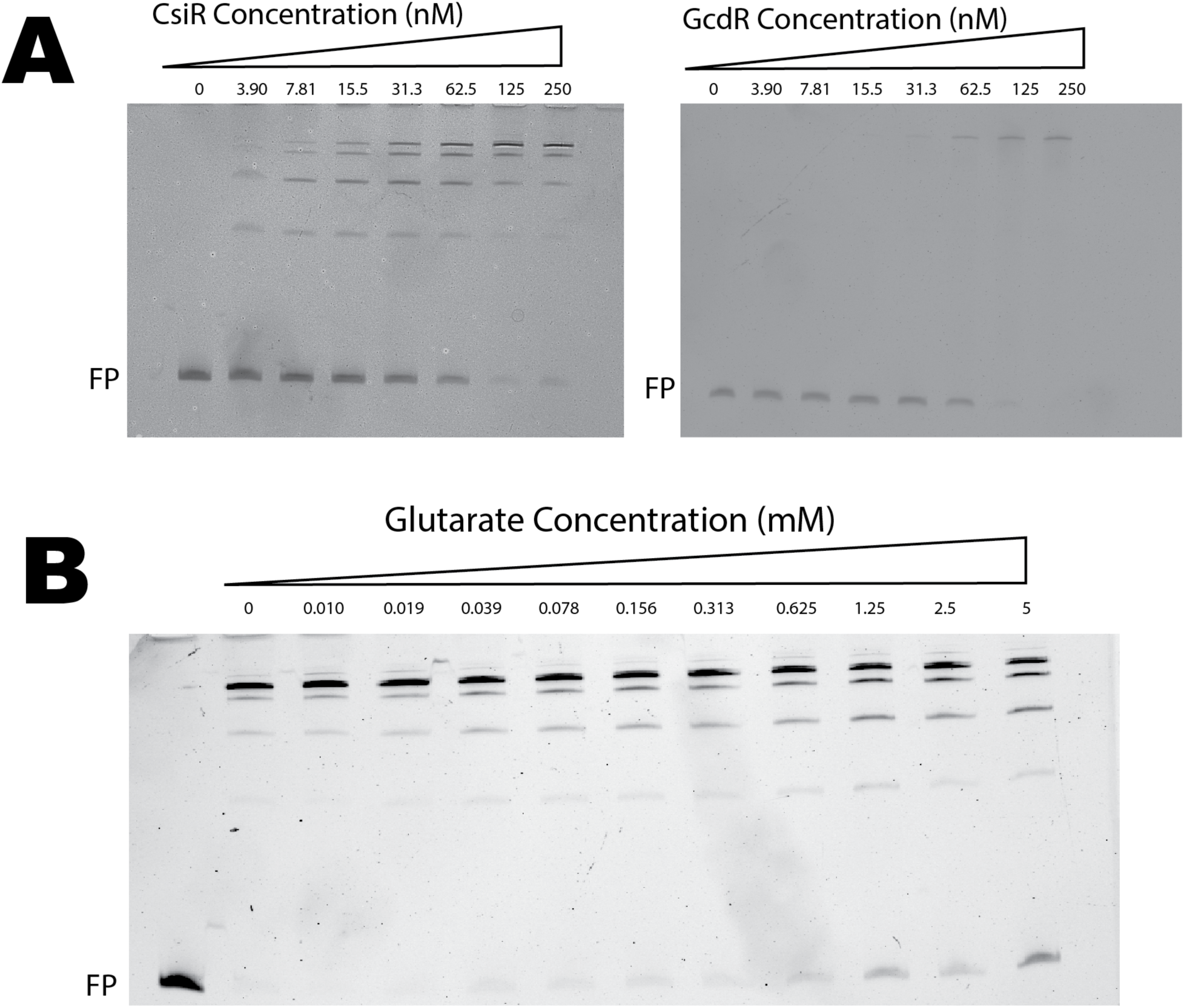
Biochemical characterization of CsiR and GcdR. (A) EMSA of either purified CsiR (left) or GcdR (right) incubated with 10 nM fluorescent probe of intergenic region between *csiR* and *csiD*. Concentrations of CsiR protein range from 0 nM to 250 nM. FP denotes free probe (B) EMSA showing 100 nM CsiR and 10 nM fluorescent probe incubated with glutarate from concentrations of 0 mM to 5 mM. Far left lane shows free probe with no CsiR. FP denotes free probe.

### Development of two glutarate biosensor vectors

In order to evaluate CsiR and GcdR as biosensors, we cloned both the regulator and intergenic region of both the coA-independent and coA-dependent glutarate catabolism pathways upstream of RFP on the broad host range vector pBADT. *E. coli* DH10B harboring either vector were grown for 24 hours in EZ-Rich medium supplemented with concentrations of glutarate ranging from 0.00015 mM to 2.5 mM, after which both OD600 and RFP fluorescence were measured. At 2.5 mM glutarate, the GcdR and CsiR vectors demonstrated a maximal induction over background of 55.5 and 1.5 times over uninduced cells, respectively; however, the CsiR system showed considerable background signal (Table 1). Normalized RFP expression for each sensor was fitted to the Hill equation to derive biosensor performance. The GcdR system was found to have a Hill coefficient of 1.33, a K_d_ of 0.32 mM, and a maximum predicted normalized RFP expression of 5403, while the CsiR system was shown to have a Hill coefficient of 1.61, a K_d_ of 0.016 mM, and a maximum predicted normalized RFP expression of 3223 (Figure 4A). To test for the ability of CsiR or GcdR to sense other diacids, *E. coli* harboring either vector were grown in LB medium with 5 mM to 0.00015 mM of either succinate, adipate, or pimelate. While pimelate and succinate were unable to induce either system, 2.5 mM adipate induced the CsiR biosensor ∼0.2x, and induced the GcdR system ∼4x over background (Figure S2).

**Figure 4:**
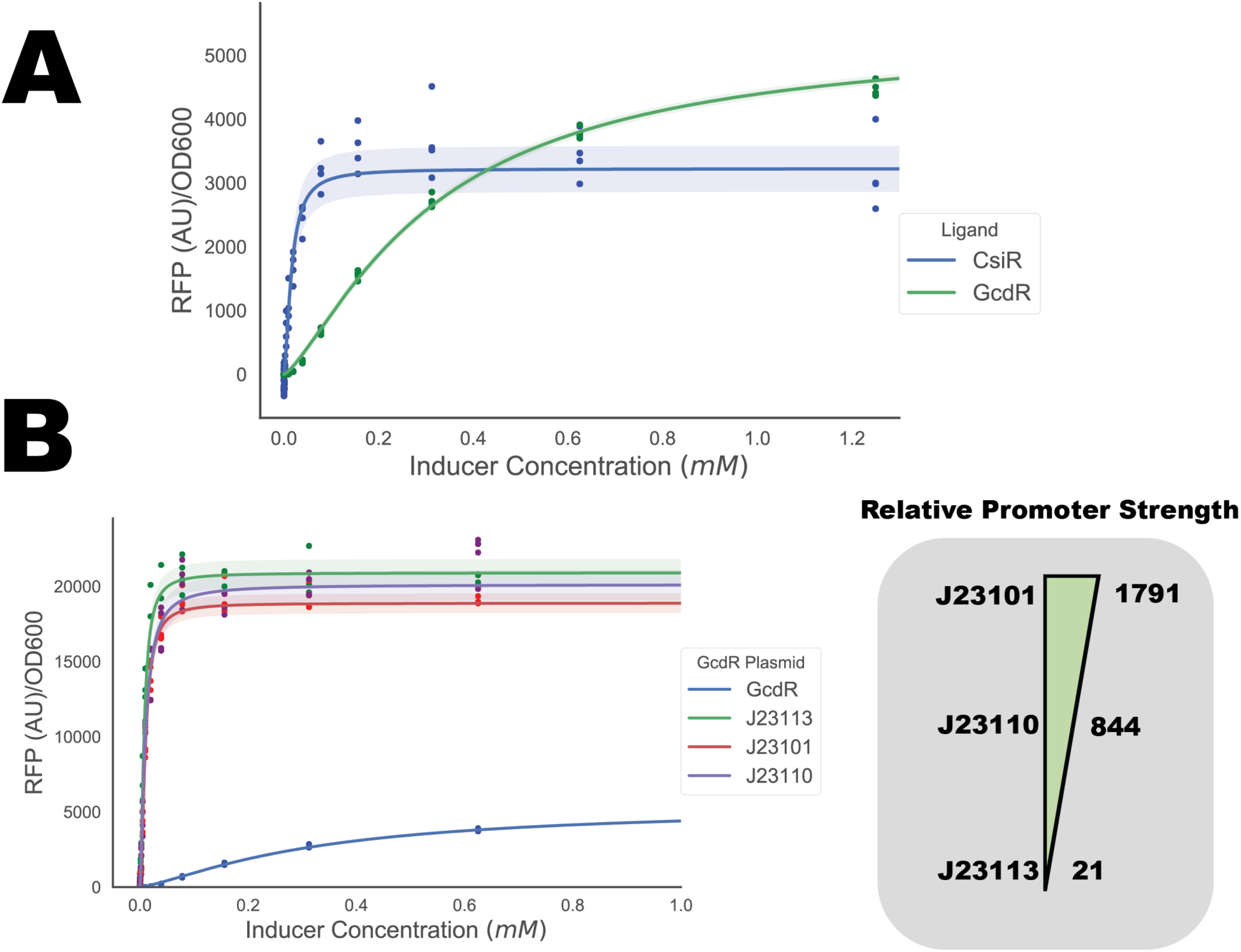
Development of glutarate responsive reporter vectors for *E. coli*. (A) Fluorescence data fit to the Hill equation to derive biosensor performance characteristics for native CsiR and GcdR systems. Points show individual experimental measurements. Shaded area represents (+/-) one standard deviation, n=4. (B) Fluorescence data fit to the Hill equation to derive biosensor performance characteristics for the engineered GcdR systems. Points show individual experimental measurements. Relative promoter strength is shown to the right. Shaded area represents (+/-) one standard deviation, n=4.

**Table 1:** Biosensor performance parameters with standard deviations. Sensitivity: Hill coefficient of fitted data. Kd: Concentration of ligand to achieve half predicted maximal RFP expression. Max (Normalized RFP): Predicted maximal RFP expression after OD normalization and subtraction of uninduced expression. Dynamic range: Minimal and maximal experimental range of OD normalized RFP. Induction: Ratio of maximal experimental induction over basal expression. Inducer Range: Experimentally determined range of glutarate that can detected with biosensor.

| Report<br>er | Sensitivi<br>ty | Kd (mM) | Max<br>(Normalized<br>RFP) | Dynamic Range<br>(RFP/OD600) | Induct<br>ion | Inducer<br>Range<br>(mM) |
| --- | --- | --- | --- | --- | --- | --- |
| CsiR-Native | 1.61 (+/-<br>0.16) | 0.016 (+/-<br>0.002) | 3223 (+/- 72) | 5704-8357 | 1.5 | 0.005-0.0195 |
| GcdR-Native | 1.33 (+/-<br>0.03) | 0.32 (+/-<br>0.006) | 5353 (+/- 38.5) | 97-5403 | 55.5 | 0.01-2.5 |
| GcdR-J23113 | 1.64<br>(+/-<br>0.09) | 0.008 (+/-<br>0.0003) | 20896 (+/-<br>184) | 3272-24027 | 7.3 | 0.001-0.0195 |
| GcdR-J23101 | 1.74 (+/-<br>0.07) | 0.01 (+/-<br>0.0003) | 18881 (+/-<br>131) | 2428-20099 | 8.3 | 0.001-0.0195 |
| **GcdR-J23110** | **1.42 (+/- 0.09)** | **0.01 (+/- 0.0004)** | **20113 (+/- 214)** | **2113-20690** | **9.8** | **0.0002-0.0195** |

**Table 2.** Strains and Plasmids used in this work

| Strain | JBEI Part ID | Reference |
| --- | --- | --- |
| *E. coli* DH10B |  | 50 |
| *E. coli* BL21(DE3) |  | Novagen |
| *E. coli* S17 |  | ATCC 47055 |
| *P. putida* KT2440 |  | ATCC 47054 |
| *P. putida* Δ*davT* | JPUB_013544 | This work |
| *P. putida* Δ*gcdH*Δ*csiD* | | 18 |
| **Plasmids** |  |  |
| pET21b |  | Novagen |
| pBADT |  | 51 |
| pMQ30 |  | 52 |
| pBADT-*gcdR*-P*gcdH*::RFP | JPUB_010960 | This work |
| pBADT-*csiR*-P*csiD*::RFP | JPUB_010962 | This work |
| pBADT-*gcdR*-J23101 | JPUB_013546 | This work |
| pBADT-*gcdR*-J23110 | JPUB_013548 | This work |
| pBADT-*gcdR*-J23113 | JPUB_013550 | This work |
| pMQ30 *davT* | JPUB_013544 | This work |
| pET21b-*CsiR*_P*put* | JPUB_010964 | This work |
| pET21b-GcdR_Pput | JPUB_010966 | This work |
| pBbS8k-rfp |  | 53 |
| pBbE8k-rfp |  | 53 |

Given that GcdR showed a greater dynamic and substrate range with no response to 2-HG, we tested whether the performance characteristics of these GcdR system could be altered. The promoter region 50 bp upstream sequence of *gcdR* in the GcdR-sensing vector was replaced by three different previously characterized constitutive promoters from the Anderson collection (J23101, J23110, and J23113), representing a high (1791 RFP AU), medium (844 RFP AU), and low (21 RFP AU) strengths ^24^. All of the engineered GcdR systems showed reduced limits of detection and increased sensitivity to glutarate compared to natively regulated GcdR, with each new vector showing a decreased K_d_ (J23113 : 0.008 mM, J23110 : 0.01 mM, J23101 : 0.01 mM) (Figure 4B).All engineered GcdR vectors showed ∼4x the maximal expression compared to the native system (Figure 4B), but also had nearly 30x greater basal RFP expression (Table 1). While all three engineered GcdR vectors performed similarly, the limit of detection of J23110 was 5x less than the other two vectors, likely due to the lower basal expression of the vector (Table 1).

### Developing metrics to quantify biosensor performance

In previously published work the mathematical basis for determining linear range or limit of detection has often been obscure and non-systematic ^1, 8, 25^. Furthermore, a linear range may not be able to adequately capture the performance characteristics of a biosensor over the range of ligand concentrations where the sensor can still resolve differences. To address these deficiencies, we sought to develop a mathematical method for evaluating the sensing performances of genetically encoded transcription factors fitted to the Hill equation. Our approach uses a probabilistic model to relate inducer concentration and corresponding fluorescence measurements fit to the Hill equation assuming: (1) fluorescence measurements at a particular concentration are normally distributed, (2) the variance of fluorescence measurements is roughly constant over the range of measured values, and (3) the relationship between ligand concentrations and fluorescence can be well modeled using the Hill function. This model allows us to estimate the concentration of ligand compatible with our observed fluorescence data given the variance of the data as determined via Markov Chain Monte Carlo (MCMC) sampling ^26^. A detailed methodological description can be found in the supplemental Jupyter notebook, which can be used to analyze other biosensor data.

By applying MCMC sampling to the model of our native GcdR biosensor, we can readily produce the probability density functions (i.e. the probability that the ligand produces the observed fluorescent response) of specific ligand concentrations (Figure 5A). At glutarate concentrations of 0.25 mM, 0.68 mM, and 1.125 mM associated fluorescence values can be resolved from one another (no overlap), however the biosensor is less able to resolve ligand concentrations between 1.5 mM and 2 mM (Figure 5A). When we apply MCMC sampling to the GcdR sensor being driven by the J23101 promoter, we observe that this system possesses the resolution to distinguish between 0.004 mM and 0.011 mM, but is less able to distinguish between higher concentrations (Figure 5B). A biosensor’s resolution window, defined as the width of the 95% prediction interval of inducer concentrations derived from a set of fluorescence measurements, can then be expressed as a continuous function across a range of ligand concentrations for a given biosensor (Figure 5C). Below concentrations of ∼0.01 mM glutarate the J23101 GcdR biosensor has greater resolution, while at higher concentrations the native GcdR sensor system has greater resolution(Figure 5C). Another important aspect of our approach is that it allows for the resolution window to be calculated as a function of the number of replicates in a biosensor experiment. If either variance decreases or sample size increases, the resolution of a biosensor also increases. By simulating sample sizes of 1 through 100 via MCMC sampling, the theoretical resolution of the native GcdR dramatically increases (Figure 4D). This “power” analysis may serve as a guide for experimental design when a certain biosensor resolution is required for a given application. We believe this approach may be generally useful to any dataset derived from fluorescent transcription factor based biosensors. To demonstrate this we also applied our MCMC methodology to two well characterized BglBrick vectors, pBbBSk-rfp and pBbBEk-rfp, which express RFP from arabinose-inducible vectors from SC101 or ColE1 origins respectively (Figure S3). Fluorescence data from each vector (Figure S3A), was fit to the Hill question (Figure S3B), and demonstrated that both vectors had the highest resolution at an arabinose concentration of ∼0.1% w/v (Figure S3C).

**Figure 5:**
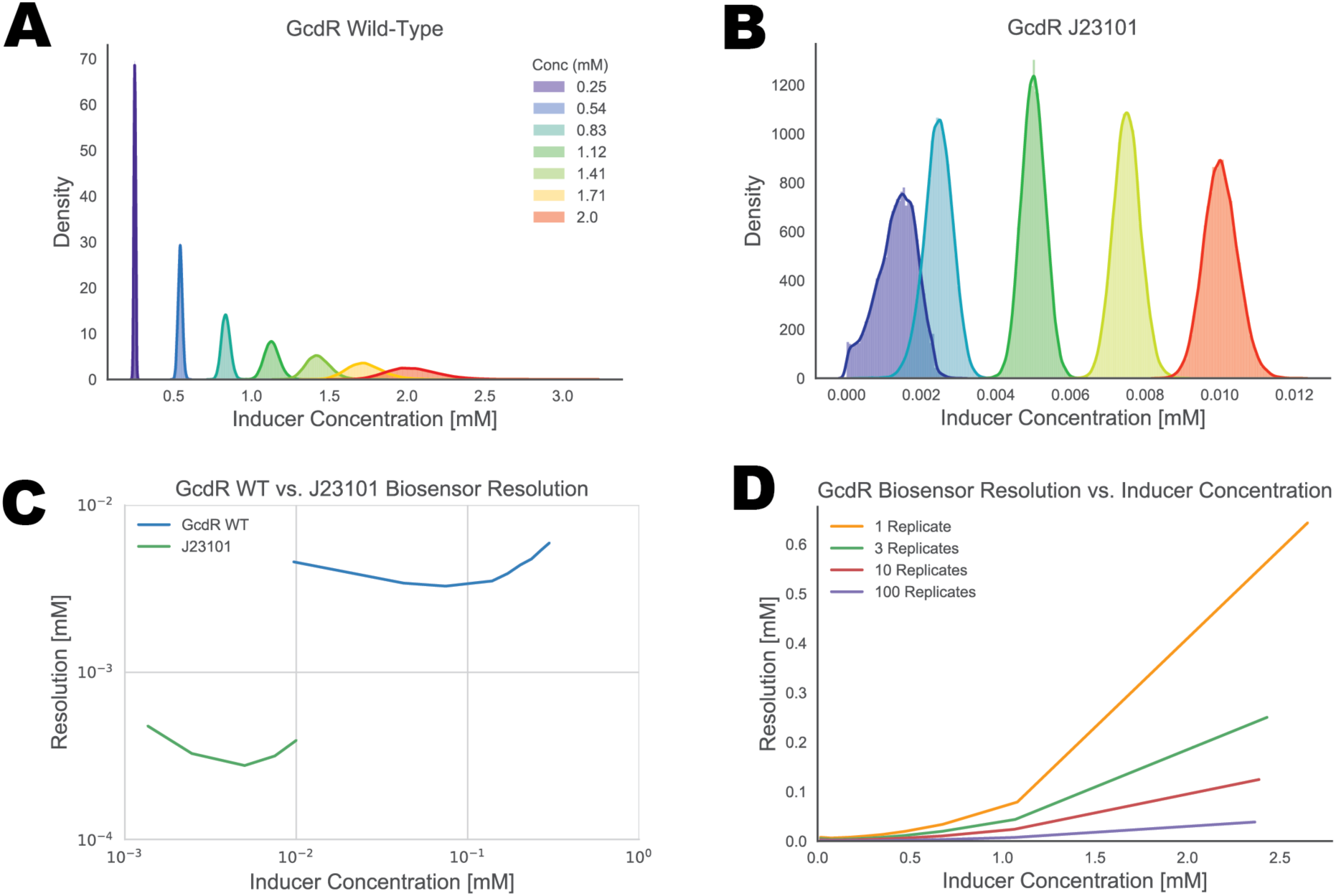
Development of analytics for biosensor performance. Probability density functions of fluorescence values compatible with concentrations of glutarate for selected ligand concentrations modelled to the Hill equation of the native GcdR system (A) or the GcdR J23101 system (B). (C) Resolution of native GcdR or GcdR J23101 biosensor systems over a select range of glutarate concentrations. (D) Theoretical resolution of the GcdR native biosensor with differing number of replicates.

### Responsiveness of glutarate biosensors to lysine metabolites in *P. putida*

To assess the ability of these vectors to function in *P. putida* both plasmids were introduced into either wild type *P. putida* or a strain with both known pathways of glutarate catabolism deleted (Δ*csiD*Δ*gcdH* - referred to as Δglutarate). The resulting strains were grown in MOPS minimal medium supplemented with 10 mM glucose and glutarate ranging from 5 mM to 0.01 mM for 24 hours. Both vectors responded to increased concentrations of exogenously applied glutarate, though the GcdR vector had ∼10x greater fluorescence than the CsiR vector (Figure 6A). The Δglutarate strain showed increased levels of RFP induction using both the CsiR and GcdR systems, suggesting both vectors are able to sense increased levels of glutarate (Figure 6A). To further examine the ability of these vectors to probe *P. putida* lysine metabolism, both were also introduced into a Δ*davT* strain, which is unable to metabolize 5-aminovalerate to glutarate semialdehyde (Figure 1A) and therefore precludes glutarate production. When wild type, Δ*davT*, and Δglutarate strains harboring either the CsiR or GcdR systems were grown on minimal medium supplemented with 10 mM glucose and 10 mM 5-aminovalerate, both vectors in the Δ*davT* strain showed decreased fluorescence compared to wild type, while vectors in the Δglutarate strains showed increased fluorescence (Figure 6B). Measurement of intracellular 5-aminovalerate showed significant pools of the metabolite in the Δ*davT* strains (∼1500 uM/OD600) with no detectable 5-aminovalerate in the other genetic backgrounds (Figure 6C). These results highly suggest that both GcdR and CsiR are insensitive to 5-aminovalerate, an essential feature of these sensors if they are to be used in organisms that derive glutarate from a 5-aminovalerate precursor.

**Figure 6:**
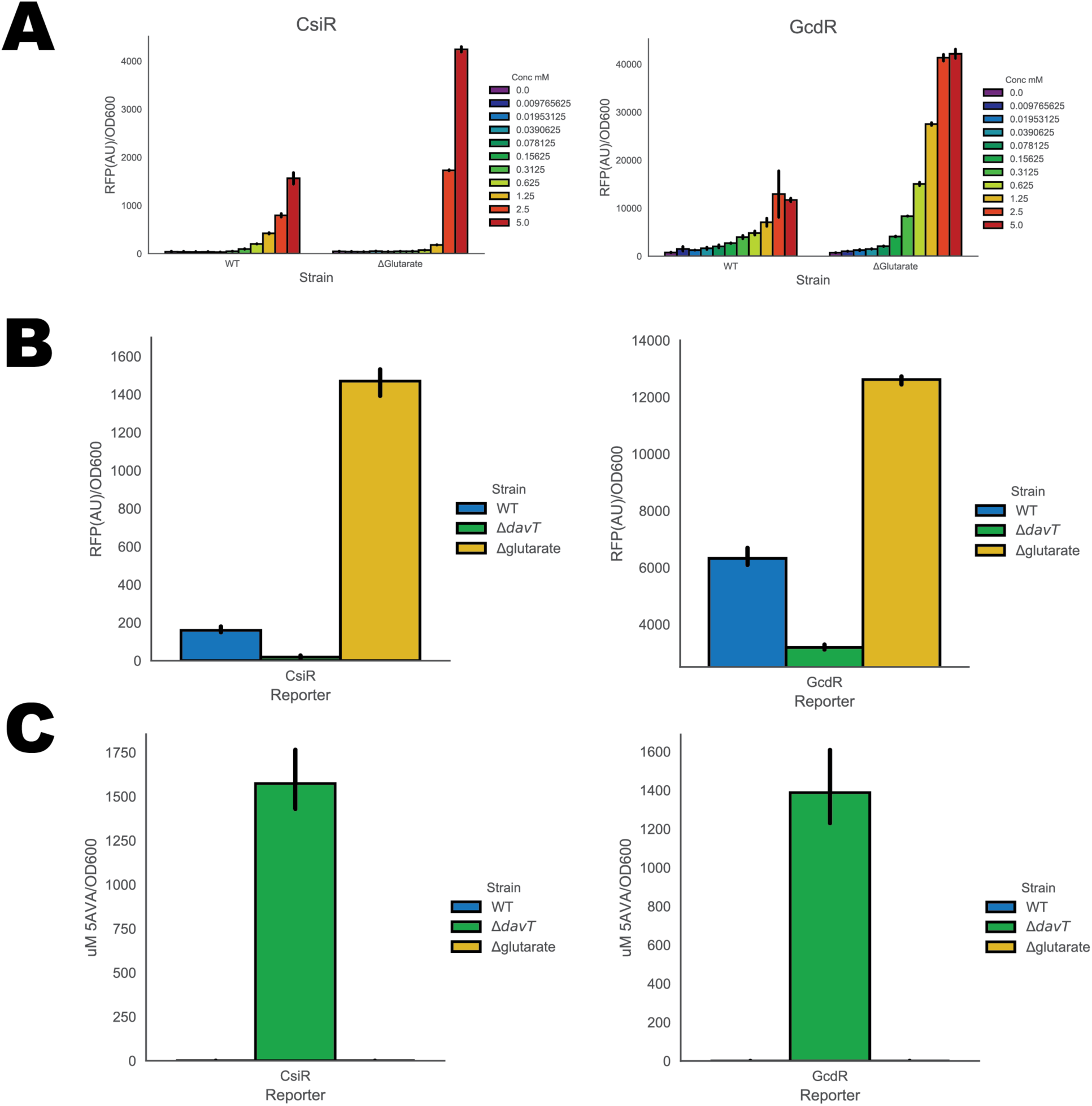
Performance of CsiR and GcdR biosensors in *P. putida*. (A) RFP expression of either wild type *P. putida* or *P. putida* with the ability to metabolize glutarate knocked out, measured with either the CsiR or GcdR biosensor under different external glutarate concentrations. Error bars represent 95% CI, n=4 (B) RFP expression of wild type, Δ*davT*, or Δglutarate strains of *P. putida* harboring either the CsiR or GcdR biosensor when grown on 10 mM glucose and 10 mM 5-aminovalerate (5AVA). Error bars represent 95% CI, n=3. (C) Intracellular concentration of 5-aminovalerate of wild type, Δ*davT*, or Δglutarate strains of *P. putida* harboring either the CsiR or GcdR biosensor when grown on 10 mM glucose and 10 mM 5-aminovalerate (5AVA). Error bars represent 95% CI, n=3.

To evaluate the ability of both reporter vectors to monitor the catabolism of other lysine metabolites, wild-type *P. putida* harboring either GcdR or CsiR were grown in minimal media supplemented with either 10 mM glucose, L-lysine, D-lysine, 5-aminovalerate, or 2-aminoadipate for 48 hours with OD_600_ and RFP fluorescence being measured continuously (Figure S4A and Figure S4B). Neither vector was induced when the bacterium was grown on glucose (Figure S4). The GcdR vector was strongly induced when grown on 5-aminovalerate, and to a lesser extent 2-aminoadipate, D-lysine, and L-lysine (Figure S4A). Conversely, the strain harboring the CsiR vector only displayed induction of RFP above background when grown on 5-aminovalerate (Figure S4B). While 5-aminovalerate was able to induce RFP induction in the strain harboring the CsiR vector, induction was extremely limited compared to induction of RFP from the GcdR vector (Figure S4).

## DISCUSSION

Recent advances in high-throughput functional genomics have allowed researchers to rapidly identify novel metabolic pathways, and in turn infer the function of novel transcription factors ^27, 28^. Rigorous characterization of these regulators is a critical step to developing novel parts for synthetic biology as well as useful tools for metabolic engineering. The discovery of two distinct pathways for glutarate catabolism within *P. putida*, regulated by independent transcription factors, presents an interesting opportunity to compare and contrast the relative sensing properties of each system.

CsiR and GcdR homologs have now been characterized thoroughly in multiple bacteria, and have demonstrated that CsiR acts as a repressor of csiD whereas GcdR acts as a positive regulator of gcdHG ^19–21^. Our bioinformatic analyses suggest that the binding site of CsiR is highly conserved amongst bacteria that possess the regulator (Figure 2). While our CsiR biosensor was shown to have a lower limit of detection of glutarate than the unengineered GcdR biosensor (Table 1), Zhang et al. demonstrated that CsiR is also responsive to 2-HG ^20^. Given that the GcdR biosensor can readily be engineered to have lower limits of detections, it is likely the better choice to detect glutarate via a genetically encoded transcription factor.

Multiple works have demonstrated that engineering changes to the expression of a transcription factor, the responsiveness of the transcriptional output can also be changed ^31, 32^. . By engineering the promoter of the *gcdR* transcription factor, the sensitivity of all the vectors were increased from ∼1.3 to ∼1.7, while the K_d_ was lowered from 0.32 mM to ∼0.01 mM (Figure 4B). This sensitivity to glutarate may make this vector useful in prototyping novel routes to biological production of the C5 diacid. However at high concentrations of adipate (>1.25 mM) GcdR is also activated, which may confound results (Figure S2).

While our preliminary work with exogenously applied ligand is promising, further work remains to be done to evaluate the ability of these sensors to detect flux of glutarate in living cells. Multiple recent publications have shown that glutarate metabolism is widespread in bacteria, and evaluating the ability of the CsiR and GcdR sensors to measure this flux will require careful experimentation with C^13^ labelled substrates. Another possible confounding factor in utilizing these vectors is the presence of CsiR or GcdR binding sites in the host organism. The overexpression of either transcription factor may misregulate host metabolism. This hypothesis is supported by the differences in growth observed between GcdR and CsiR biosensor containing *P. putida* when grown on lysine metabolites (Figure S4).

The overarching goal of synthetic biology is to apply engineering principles to biological systems so that outcomes of genetic manipulation can become more predictable and repeatable ^33, 34^. While there has been a large body of work devoted to the characterization and application of biosensors, there has been conspicuously few attempts to rigorously describe at which ligand concentrations a biosensor is useful. Often the analysis merely states limit of detection, and a ‘linear range’ with little in terms of a mathematical justification. Here we present an alternative metric that allows for the calculation of ligand resolution across the entire range of detection for a biosensor. By leveraging a MCMC approach to predicting ligand ranges compatible with fluorescence values, researchers can more precisely describe a biosensors performance and identify whether a given biosensing system is potentially useful for a given engineering task. The MCMC approach also allows for the simulation of an increasing number of replicates, which could inform the researcher of the replicates that may be required in an experimental design to achieve a desired level of resolution. We hope that this initial work to better characterize biosensor performance inspires other groups to develop even more sophisticated methods of analysis.

In addition to their utility as biosensors for metabolic engineering, these sensors may be a valuable tool in studying the carbon utilization in the native host *P. putida*. Work conducted here demonstrates the ability of both CsiR and GcdR sensors to distinguish between glutarate accumulating and mutants blocked in their ability to metabolize 5-aminovalerate to glutarate (Figure 6). Lysine metabolism in *P. putida* is isomer specific, with each isomer being degraded by a separate catabolic pathway ^18^. While cross-feeding between the pathways has been proposed previously ^35^, recent work by our group has proposed a molecular mechanism for metabolite exchange between the D- and L-catabolic pathways ^18^. The exchange relies on the 2-oxoacid promiscuity of the non-heme Fe(II) oxidase CsiD, which normally catalyzes the hydroxylation of glutarate using 2-ketoglutarate as a cosubstrate to yield 2-hydroxyglutarate and succinate. CsiD can also use 2-oxoadipate, a D-lysine catabolic intermediate, as a 2-oxoacid cosubstrate to yield 2-hydroxyglutarate and glutarate as products. The glutarate from this reaction could then proceed down the L-lysine catabolic pathway. When 2-aminoadipate was fed into *P. putida* harboring the GcdR vector, fluorescence was observed in stationary phase. As 2-aminoadipate immediately precedes 2-oxoadipate in the D-lysine catabolic pathway, these results support the hypothesis that CsiD could act as a bridge between the two catabolic pathways. There has been substantial interest in developing microbes to produce glutarate, with strains of *E. coli*, *P. putida*, and *Corynebacterium glutamicum* all engineered to produce high titers ^16, 36, 37^. Further engineering of the GcdR system may be able to extend the resolvable range to higher concentrations, furthering its utility as a tool to achieve even higher titers of glutarate. Though glutarate is a valuable commodity chemical, the C6 diacid adipate is used in much greater quantities primarily as a monomer used to make nylons 33. This slight sensitivity of GcdR toward adipic acid is especially interesting, as recent work has demonstrated the effectiveness of evolving transcription factors to sense non-native ligands ^6^. Such methods could be applied to GcdR in order to expand its utility in sensing other industrially important diacids.

## METHODS

### Media, chemicals, and culture conditions

*E. coli* cultures were grown in Luria-Bertani (LB) Miller medium (BD Biosciences, USA) at 37 °C while *P. putida* was grown at 30 °C. When indicated, *P. putida* and *E. coli* were grown on modified MOPS minimal medium ^39^. Cultures were supplemented with kanamycin (50 mg/L, Sigma Aldrich, USA), gentamicin (30 mg/L, Fisher Scientific, USA), or carbenicillin (100 mg/L, Sigma Aldrich, USA), when indicated. All other compounds were purchased through Sigma Aldrich (Sigma Aldrich, USA) .

### Strains and plasmids

All bacterial strains and plasmids used in this work are listed in Table 3. All strains and plasmids created in this work are available through the public instance of the JBEI registry. (https://public-registry.jbei.org/folders/390). All plasmids were designed using DeviceEditor and VectorEditor software, while all primers used for the construction of plasmids were designed using j5 software ^40–42^. Plasmids were assembled via Gibson Assembly using standard protocols ^43^, or Golden Gate Assembly using standard protocols ^44^. Plasmids were routinely isolated using the Qiaprep Spin Miniprep kit (Qiagen, USA), and all primers were purchased from Integrated DNA Technologies (IDT, Coralville, IA). Construction of *P. putida* deletion mutants was performed as described previously ^18^.

### Expression and purification of proteins

Proteins were purified as described previously ^45^. Briefly, 500 mL cultures of *E. coli* BL21 (DE3) harboring expression plasmids were grown in LB medium at 37 °C to an OD of 0.6 then induced with 1mM isopropyl β-D-1-thiogalactopyranoside. Cells were allowed to express for 18 hours at 30 °C before being harvested via centrifugation. Cell pellets were stored at -80 °C until purification. Cell pellets were then resuspended in lysis buffer (50 mM sodium phosphate, 300 mM sodium chloride, 10 mM imidazole, 8% glycerol, pH 7.5) and sonicated to lyse cells. Insolubles were pelleted via centrifugation (30 minutes at 40,000 x g). The supernatant was applied to a fritted column containing Ni-NTA resin (Qiagen, USA) which had been pre-equilibrated with several column volumes of lysis buffer. The resin was washed with lysis buffer containing 50 mM imidazole, then the protein was eluted using a stepwise gradient of lysis buffer containing increasing imidazole concentrations (100 mM, 200 mM, and 400 mM). Fractions were collected and analyzed via SDS-PAGE. Purified proteins were concentrated using Spin-X UF 20 (10 kDa MWCO) spin concentrators (Corning, Inc.). Concentrated protein was stored at 4 °C until *in vitro* analysis.

### Plate based growth and fluorescence assays in P. putida

Growth studies of bacterial strains were conducted via microplate reader kinetic assays. Overnight cultures were inoculated into 10 mL of LB medium from single colonies, and grown at 30 °C. These cultures were then washed twice with MOPS minimal medium without any added carbon and diluted 1:100 into 500 uL of MOPS medium with 10 mM of a carbon source in 48-well plates (Falcon, 353072). Plates were sealed with a gas-permeable microplate adhesive film (VWR, USA), and then optical density and fluorescence were monitored for 48 hours in a Biotek Synergy 4 plate reader (BioTek, USA) at 30 °C with fast continuous shaking. Optical density was measured at 600 nm, while fluorescence was measured using an excitation wavelength of 485 nm and an emission wavelength of 620 nm with a manually set gain of 100.

### Transcriptional fusion fluorescence assays

To measure RFP production in*E. coli,* fluorescence measurements were obtained from single time points of cells grown in deep-well 96-well plates as described previously with minor changes^46^. Briefly, cells were grown in 500 µL of EZ-Rich medium supplemented with kanamycin and a range of glutarate concentrations from 5 mM to 0 mM. Plates were sealed with AeraSeal film (Excel Scientific, AC1201-02) and grown for 22 hours at 37 °C on a 200 rpm shaker rack. After incubation, 100 µL from each well was aliquoted into a black, clear-bottom 96-well plate and fluorescence was measured with a Tecan Infinite F200 plate reader (Tecan, USA). Optical density was measured at 600 nm, while fluorescence was measured using an excitation wavelength of 535 nm and an emission wavelength of 620 nm with a manually set gain of 60. Endpoint fluorescence assays in P. putida were carried out in LB media, and measured by the same method.

### Electrophoretic mobility shift assays

Electrophoretic mobility shift assays were performed as previously described ^7^. 6-Carboxyfluorescein labelled PCR products for CsiR probes were generated from the intergenic region between PP_2908 and PP_2909 using primers csiRprobeFOR 5’-6-FAM/AGTTCGATCTGCGTAAAG-3’ and csiRprobeREV 5’-CCCGCTGAATGCTGAGTT-3’), while probes for GcdR were generated from the intergenic region between PP_0157 and PP_0158 with primers gcdRprobeFOR 5’-6-FAM/CGGGTCGATCCAGTTGAAA-3’ and gcdRprobeREV 5’-GCATGTACGTCAACCTCACT-3’. Primers were purchased from IDT Technologies (IDT, Coralville, IA). PCR product was then purified with a QIAquick PCR Purification Kit (Qiagen, USA), and the amount of DNA was quantified via a NanoDrop 2000C (Thermo Fisher Scientific). Binding reactions were conducted with 10 ng of labelled probe, which was added to 10 mM Tris–HCl (pH 8.0), 25 mM KCl, 2.5 mM MgCl2, 1.0 mM DTT and 2 ug salmon sperm DNA in 20 uL reactions. CsiR was added to reactions in concentrations ranging from 250 nM to ∼4 nM, in addition to a control reaction without CsiR, and then allowed to incubate at 22 °C for 20 minutes. Reactions were then loaded into 10% native polyacrylamide gels buffered with 0.5x TBE. Afterwards, electrophoresis gels were imaged on an Alpha Innotech MultiImage III (Alpha Innotech). Adobe Photoshop was used to average pixel intensity over the entire band on EMSA gels in order to estimate the K_d_.

### Measurement of 5-aminovalerate

To measure intracellular concentrations of 5-aminovalerate, cells were quenched as previously described ^47^. LC/MS analysis was performed on an Agilent 6120 single quadrupole LC/MS equipped with a Waters Atlantis Hilic 5 µM Silica column (4.6 x 150 mm). A linear gradient of 100-30% 90:10 CH3CN:H20 with 10 mM ammonium formate and 0.1% formic acid (v/v) over 20 min in 90:10 H2O:CH3CN with 10 mM ammonium formate and 0.1% formic acid (v/v) at a flow rate of 1.0 mL/min was used. Extracted ion chromatograms were integrated and peak area was used to construct a standard curve using an authentic 5-aminovalerate standard. Concentrations of 5-aminovalerate within samples were interpolated from this curve.

### Analysis of biosensor parameters

A model relating inducer concentrations and fluorescence measurements to characterize the performance of a biosensor was generated under the following assumptions 1) the relationship between analyte concentrations and fluorescence can be well modeled using the Hill equation 2) fluorescence measurements at a particular concentration are normally distributed 3) the variance of fluorescence measurements is roughly constant over the range of measured values.

Under these assumptions we can phrase the following probabilistic model via equation 1 (Figure 7). Using the probabilistic model which captures our constraints on the problem the log likelihood function is expressed as equation 2 (Figure 7).The log likelihood is used to express the maximum likelihood estimation (MLE) problem as equation 3 (Figure 7),which when solved results in the optimal parameters of the model given the characterization data. In order to estimate the distribution of ligand concentrations that are compatible with experimental fluorescence data, MCMC sampling was used to solve the MLE problem equation 4 (Figure 7). We determined biosensor resolution by solving the above maximum likelihood estimation problem iteratively over the range of observed fluorescences during the biosensor characterization process. This can determine the relationship between an inducer concentration estimate and the estimated standard deviation. The standard deviation of the estimate of inducer concentration can be interpreted as the resolution window. In the case of this work, two standard deviations is considered the resolution window of the sensor, as 95% of the compatible inducer concentration estimates fall within the interval.

**Figure 7:**
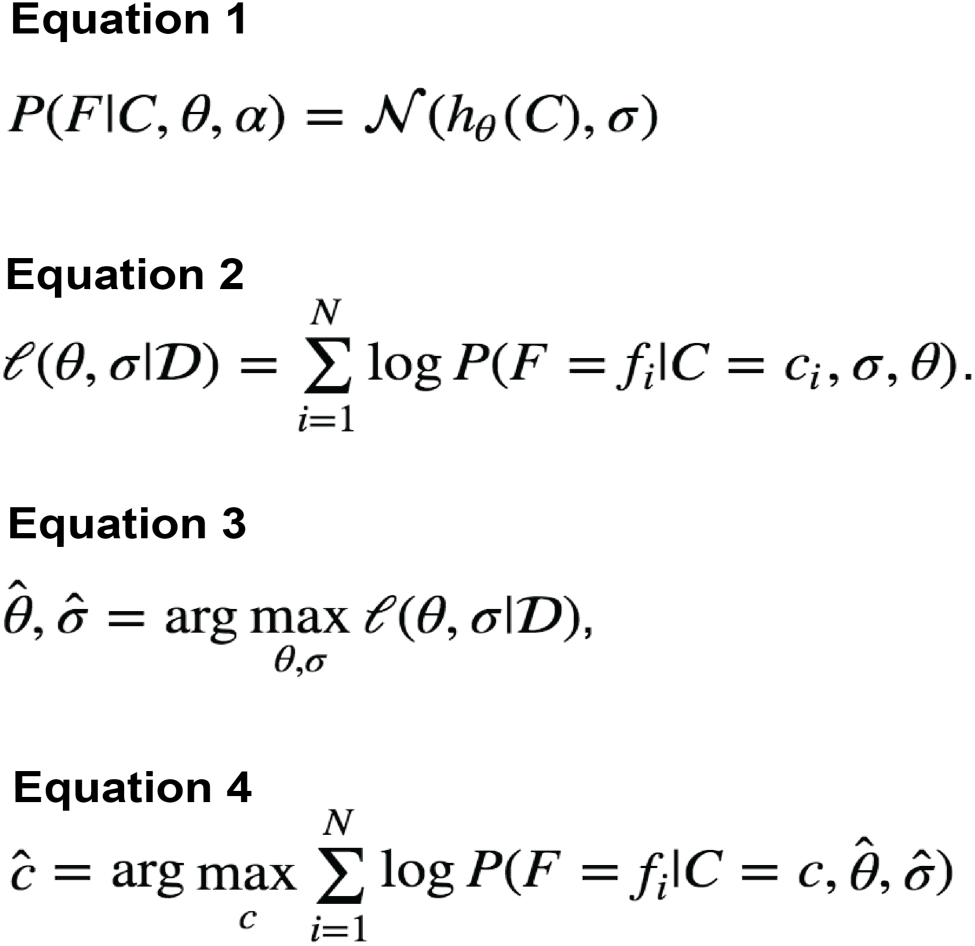
Equations used in this study for MCMC analysis of biosensor data.

To determine minimal and maximal levels of ligand detection of a given biosensor, the minimal detection limit was defined as the minimal concentration of inducer that resulted in a OD600 normalized RFP value significantly (t-test pval <0.05) greater than that of uninduced cultures, while the maximal detection limit was defined as the greatest concentration of inducer that resulted in a OD600 normalized RFP value significantly (t-test pval <0.05) less than that of cultures induced with the highest concentration of ligand experimentally tested (2.5 mM).

A comprehensive methodological description of calculating biosensor performance parameters, as well as usable Jupyter notebooks can be found at https://github.com/JBEI/biosensor_characterization_public.

### Bioinformatic analysis

For the phylogenetic reconstructions, the best amino acid substitution model was selected using ModelFinder as implemented on IQ-tree ^48^, phylogenetic trees were constructed using IQ-tree, and nodes were supported with 10,000 bootstrap replicates. The final tree figures were edited using FigTree v1.4.3 (http://tree.bio.ed.ac.uk/software/figtree/). Orthologous syntenic regions of CsiR were identified with CORASON-BGC ^49^ and manually colored and annotated. DNA-binding sites were predicted with MEME ^22^.

## SUPPORTING INFORMATION

The Supporting Information is available free of charge on the ACS Publications website.

S1.pdf contains supplementary figures.

## Acknowledgements

The authors would like to thank Tyler Backman for assistance creating Git repositories. The authors would like to thank the UC Berkeley SMART program for providing support for R.K to conduct summer research. This work was part of the DOE Joint BioEnergy Institute (https://www.jbei.org) supported by the U. S. Department of Energy, Office of Science, Office of Biological and Environmental Research, and was part of the Agile BioFoundry (http://agilebiofoundry.org) supported by the U.S. Department of Energy, Energy Efficiency and Renewable Energy, Bioenergy Technologies Office, through contract DE-AC02-05CH11231 between Lawrence Berkeley National Laboratory and the U.S. Department of Energy. H.G.M. was also supported by the Basque Government through the BERC 2018-2021 program and by Spanish Ministry of Economy and Competitiveness MINECO: BCAM Severo Ochoa excellence accreditation SEV-2017-0718.The views and opinions of the authors expressed herein do not necessarily state or reflect those of the United States Government or any agency thereof. Neither the United States Government nor any agency thereof, nor any of their employees, makes any warranty, expressed or implied, or assumes any legal liability or responsibility for the accuracy, completeness, or usefulness of any information, apparatus, product, or process disclosed, or represents that its use would not infringe privately owned rights. The United States Government retains and the publisher, by accepting the article for publication, acknowledges that the United States Government retains a nonexclusive, paid-up, irrevocable, worldwide license to publish or reproduce the published form of this manuscript, or allow others to do so, for United States Government purposes. The Department of Energy will provide public access to these results of federally sponsored research in accordance with the DOE Public Access Plan (http://energy.gov/downloads/doe-public-access-plan).

## Contributions

Conceptualization, M.G.T.; Methodology, M.G.T.,Z.C, J.M.B, P.C.M.,N.H.,W.S. Investigation, M.G.T.,Z.C., J.M.B, R.N.K, P.C.M, M.R.I.,A.N.P..; Writing – Original Draft, M.G.T, M.R.I..; Writing – Review and Editing, All authors.; Resources and supervision, J.D.K,P.S.,H.M.

---

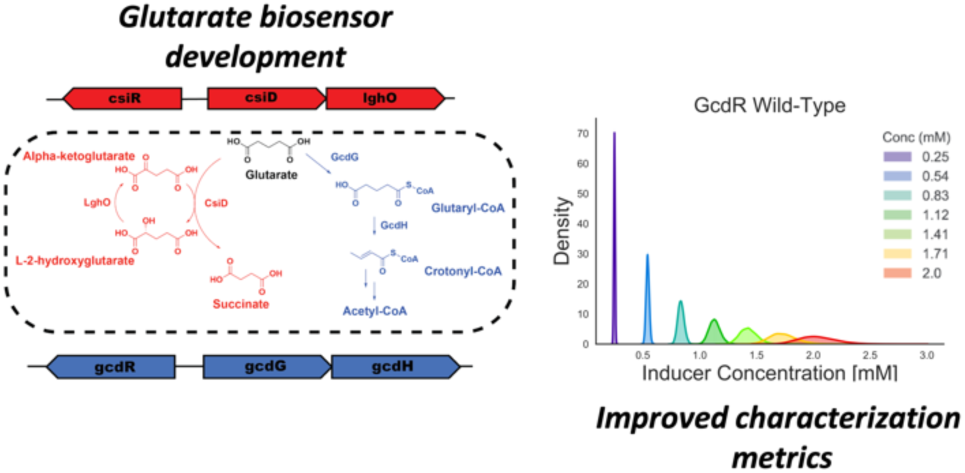

