## Supplementary Figures for "Robust characterization of two distinct glutarate sensing transcription factors of *Pseudomonas putida* L-lysine metabolism"

### Supporting Information

### Supplementary Figures

Figure S1. SDS-PAGE gels showing the purification of GcdR (A), and CsiR (B) via affinity purification.

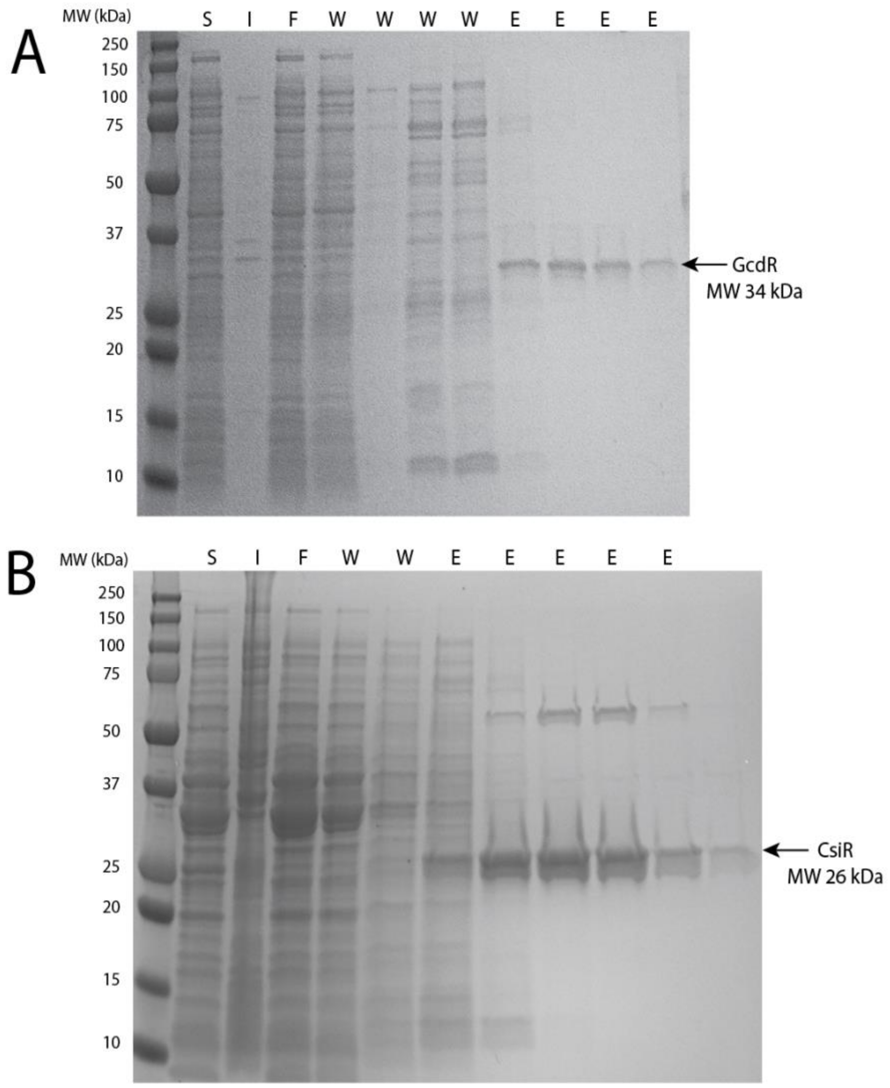


S = Soluble
I = Insoluble
F = Flowthrough
W = Wash
E = Elution

**Figure S2. Induction of the CsiR (A), or GcdR (B) biosensor vectors by succinate, adipate, and pimelate. Error bars represent 95% CI, n=4.**

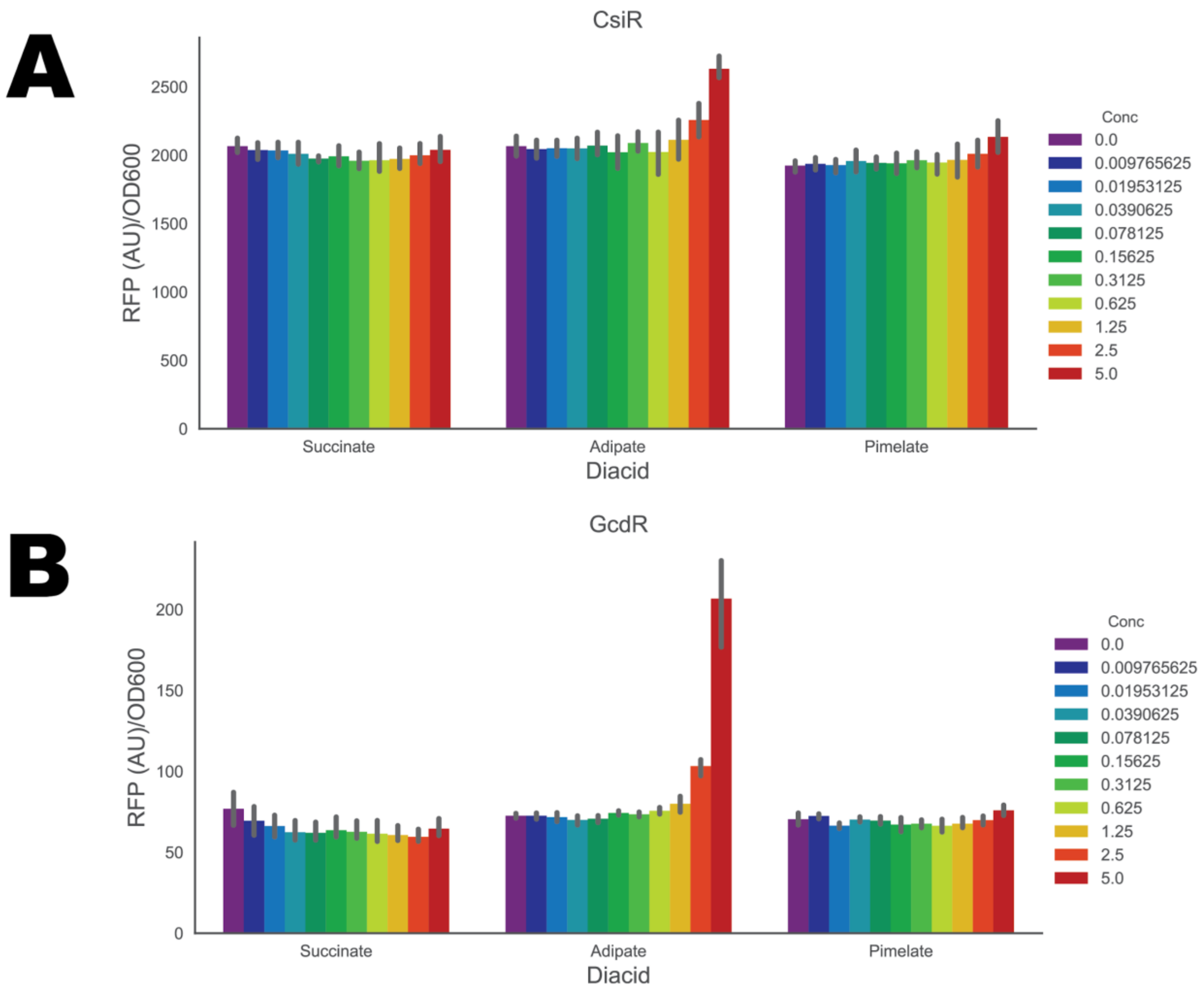

**Figures S3. Application of MCMC methodology to characterized BglBrick vectors. (A) Raw RFP expression data from SC101 and ColE1 arabinose inducible BglBrick vectors with increasing concentrations of arabinose (%w/v). (B) OD600 normalized and basal-expression zeroed fluorescence data fit to the Hill equation. Points show individual experimental measurements. n=4 (C) MCMC predicted resolution of SC101 or ColE1 biosensor systems over a select range of arabinose concentrations %w/v.**

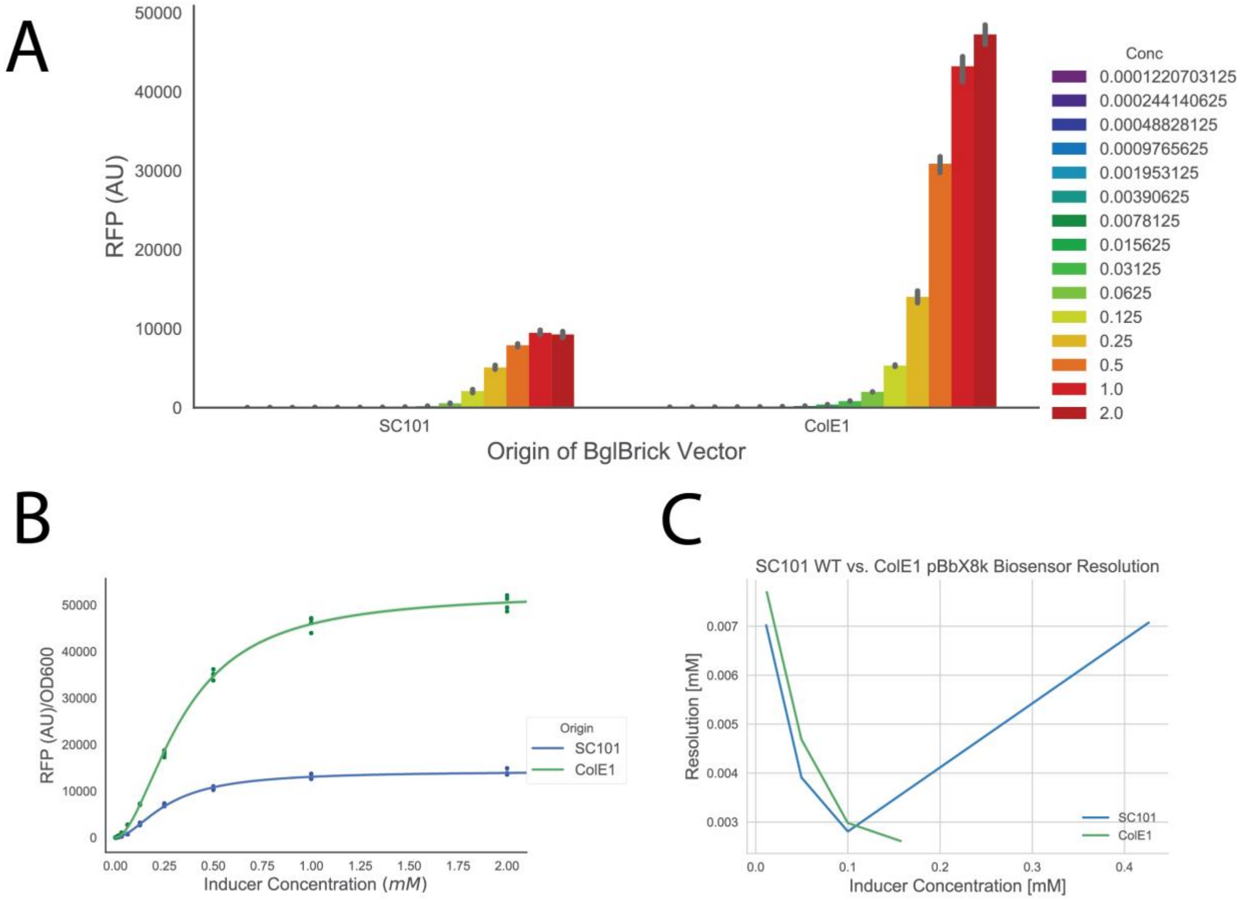

**Figures S4. Growth and fluorescence of *csiR* and *gcdR* vectors in P. putida KT2440 (A) Growth of strains harboring *gcdR* vector (left), RFP fluorescence (right) on MOPS minimal media supplemented with 10mM of glucose, L-lysine, D-lysine, 5-aminovalerate, or 2-aminoadipate as a sole carbon source. Shaded region represents 95% CI, n=3. (B) Growth of strains harboring *csiR* vector (left), RFP fluorescence (right) on MOPS minimal medium supplemented with 10mM of glucose, L-lysine, D-lysine, 5-aminovalerate, or 2-aminoadipate as a sole carbon source. Shaded region represents 95% CI, n=3.**

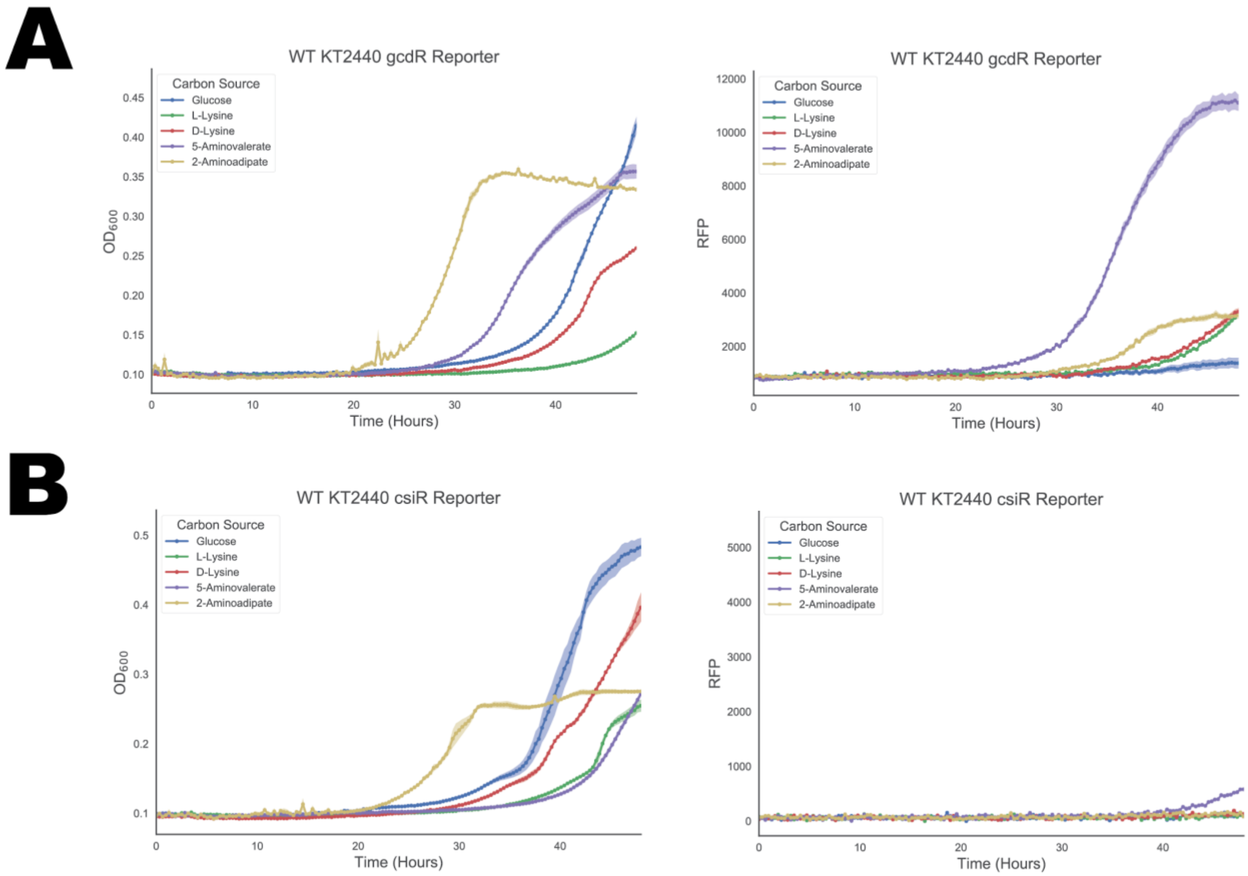
